# Loss of retinogeniculate synaptic function in the DBA/2J mouse model of glaucoma

**DOI:** 10.1101/2022.08.15.503974

**Authors:** Jennie C. Smith, Kevin Yang Zhang, Asia Sladek, Jennifer Thompson, Elizabeth R. Bierlein, Ashish Bhandari, Matthew J. Van Hook

## Abstract

**Background:** Retinal ganglion cell (RGC) axons comprise the optic nerve and carry information to the dorsolateral geniculate nucleus (dLGN) that is relayed to the cortex for conscious vision. Glaucoma is a blinding neurodegenerative disease that commonly results from intraocular pressure (IOP)-associated injury leading to RGC axonal pathology, disruption of RGC outputs to the brain, and eventual apoptotic loss of RGC somata. The consequences of elevated IOP and glaucomatous pathology on RGC signaling to the dLGN are largely unknown and likely to be important contributors to visual system dysfunction in glaucoma. Thus, the goal of this study was to determine how glaucoma affects RGC outputs to the dLGN.

**Methods:** We used a combination of anatomical and physiological approaches to study the structure and function of retinogeniculate synapses in male and female DBA/2J mice at multiple ages before and after IOP elevation. These included measures of anterograde axonal transport, immunofluorescence staining of RGC axon terminals, patch-clamp recording retinogeniculate (RG) synapses in living brain slices, Sholl analysis of thalamocortical relay neuron dendrites, measurements of RGC somatic density, and treatment with a topical ophthalmic alpha-2 adrenergic agonist (brimonidine).

**Results:** DBA/2J mice showed progressive loss of anterograde optic tract transport to the dLGN and vGlut2 labeling of RGC axon terminals. Patch-clamp measurements of RG synaptic function showed that the strength of synaptic transmission was lower in 9 and 12-month DBA/2J mice and that this was the result of loss of individual RGC axon contributions. TC neuron dendrites showed a reduction in complexity at 12 months, suggestive of a delayed reorganization following reduced synaptic input. There was no detectable change in RGC soma density in 11-12m DBA/2J retinas indicating that observed effects occurred prior to RGC somatic loss. Finally, treatment with brimonidine eye drops prevented the loss of vGlut2-labeled RGC terminals in the dLGN.

**Conclusions:** These findings identify glaucoma- and IOP-associated functional deficits in an important subcortical RGC projection target. This sheds light on the processes linking IOP to vision loss and will be critical for informing future diagnostic approaches and vision-restoration therapies.

## Introduction

Glaucoma is a neurodegenerative disease commonly associated with a sensitivity to intraocular pressure (IOP) and progressive degeneration of retinal ganglion cells (RGCs), the output neurons of the retina [1–3]. The goal of this study was to determine the timing and mechanisms by which IOP leads to loss of RGC output synapses (retinogeniculate/RG synapses) in the dorsolateral geniculate nucleus (dLGN), a subcortical RGC retinal projection target in the thalamus where convergent RGC synaptic inputs to thalamocortical (TC) relay neurons drive TC neuron action potential output to the visual cortex for conscious, “image-forming” vision. Perturbations to dLGN function during glaucoma are likely to contribute to vision loss, yet the functional impacts of elevated IOP and glaucoma on dLGN function are poorly understood.

The mechanisms of visual impairment in glaucoma are commonly viewed through the lens of dysfunction progressing toward late-stage apoptotic loss of RGCs. This process occurs as the result of an IOP- and age-induced injury to RGC axons at the optic nerve head [4], where the axons exit the eye, triggering retrograde effects on RGCs and their presynaptic partners in the retina. This involves early remodeling of RGC dendrites [5–14], alterations in their intrinsic excitability and response properties [5–7,14], and reorganization of bipolar cell ribbon synapses [6,7,15,16]. However, elevated IOP also has effects that profoundly alter the function of RGC axons distal to the optic nerve head as well as to their downstream visual targets in the brain. These include disruption of optic nerve active transport [17], metabolism [18–21], and glia [22] as well as alterations to mitochondria [23], RGC excitatory output synapses [17,24], and the structure and response properties of neurons residing in visual brain nuclei [14,25]. Evidence to date indicates that many of these functional changes are relatively early events in the pathological process. We have shown previously that elevated IOP and optic nerve injury lead to changes in the strength of retinogeniculate (RG) synaptic transmission and to alterations in TC neuron dendritic structure and intrinsic excitability [25,26]. Evidence from primate and human studies indicates that glaucoma leads to dendritic remodeling and neuronal atrophy within the visual thalamus [27–30]. In the superior colliculus of DBA/2J mice with inherited glaucoma, ultrastructural studies show that glaucoma leads to atrophy of presynaptic RGC axon terminals, reduced mitochondrial volume, and decreased size of presynaptic active zones [24].

While prior findings suggets that glaucoma can alter the function of retinal output synapses in the brain, the nature of these functional deficits, their time course, and the underlying mechanisms are largely unknown. Any disruption of visual information flow from the retina to the brain is likely to make an early contribution to visual deficits triggered by increased IOP. Moreover, understanding the timing of events triggered by IOP elevation will be critical in designing approaches to assess vision loss and interventions to halt disease progression and re-establish retinal connections with central visual targets through axon regeneration and/or stem cell replacement approaches.

Here, we made use of the DBA/2J mouse model of glaucoma [31–33], a commonly used model system that recapitulates many key features of human glaucomatous neurodegeneration, to probe how IOP elevation leads to the loss of RGC output synapses in the dLGN. Using a combination of experimental approaches to assess optic nerve transport, RGC survival, and RG synaptic structure and function, we find that DBA/2J mice show IOP-dependent deficits in RGC axonal function that are followed by progressive loss of vGlut2-labeled RGC axon terminals in the dLGN and progressive loss of functional RGC synaptic inputs to each TC neuron. This is accompanied by late-stage reorganization of TC neuron dendrites. Notably, these deficits appear to occur prior to major RGC cell body loss, as assessed using immunofluorescence staining in retinal flat mounts. Finally, we find that treatment with an ophthalmic alpha-2 receptor agonist (brimonidine eye drops), a medication used to lower IOP in glaucoma patients that also has documented neuroprotective effects [34–37], preserves vGlut2-labeled RGC axon terminals in the dLGN. Thus, we establish the functional consequences of elevated IOP that lead to the loss of conveyance of visual signals from RGCs to their post-synaptic targets in the dLGN. The loss of functional RGC output synapses is a major feature of neurodegenerative disease progression and likely to contribute to glaucomatous vision loss.

## Materials and Methods

### Animals

Animal protocols were approved by the Institutional Animal Care and Use Committee at the University of Nebraska Medical Center. Male and female DBA/2J (D2, Jackson Labs #000671, RRID:IMSR_JAX:000671) and DBA/2J-gpnmb+ (D2-control, Jackson Labs #007048, RRID:IMSR_JAX:007048) [33,38] were bred in-house and housed on a 12h/12h light/dark cycle with standard food and water. Intraocular pressure (IOP) was measured approximately monthly beginning at 2 months of age using an iCare Tonolab rebound tonometer (iCare, Vantaa, Finland) in mice that were lightly anesthetized with isoflurane. Measurements were taken within 3 minutes of isoflurane anesthesia to minimize effects of the anesthesia on IOP. Mice were euthanized by inhalation of CO_2_ followed by cervical dislocation, in keeping with American Veterinary Medical Association guidelines on euthanasia. A subset of mice was treated with Brimonidine tartrate eye drops (0.2%), one drop per eye 4-5 days per week from approximately 6-9 months of age. A control group of mice received treatment with lubricating eye drops lacking brimonidine (Systane, Alcon).

### Cholera toxin beta injections and analysis

To test for deficits in anterograde transport along the optic tract, mice received a unilateral injection of cholera toxin beta subunit coupled to Alexa Fluor 594 (CTb-594, Invitrogen C34777). Mice were anesthetized with isoflurane and treated with proparacaine ophthalmic drops (1%). A Hamilton syringe and 33 gauge needle were used to deliver a unilateral intravitreal injection of ^~^1-2 μL of CTb-594 (1 μg/mL). 3-4 days post-injection, mice were euthanized with CO_2_ asphyxiation and cervical dislocation. Brains were dissected, rinsed briefly in phosphate buffered saline (PBS), and fixed by immersion in 4% paraformaldehyde in PBS for overnight. After fixation, brains were rinsed in PBS, cryoprotected overnight in 30% sucrose, embedded in 3% agar, and sliced into 100 micron-thick slices on a Leica VT1000S vibratome. Every other section containing the dLGN was mounted on SuperFrost Plus slides (Fisher Scientific) and coverslipped with Vectashield Hardset (Vector). CTb-594 images of the contralateral dLGN were acquired using a 10x objective lens on an Olympus BX51WI microscope with a Tucsen monochrome camera. To analyze CTb-594 labeling, each image was thresholded in ImageJ based on a region outside of the dLGN and the number of CTb-594 pixels was counted using the histogram. In this way, the total volume of the dLGN labeled by CTb-594 was calculated in serial dLGN sections for each mouse.

### Immunofluorescence staining

Retinal ganglion cell axon terminals were labeled by immunofluorescence staining for vGlut2. After euthanasia, brains were dissected into PBS and fixed for 4h in 4% PFA. Brains were then rinsed in PBS, cryoprotected overnight in 30% sucrose, embedded in 3% agar, and cut into 50 micron-thick sections using a Leica VT1000S vibratome. Sections were mounted on SuperFrost Plus slides. For staining, sections were rinsed in PBS, blocked/permeabilized with 0.5% TritonX-100, 5.5% goat serum and 5.5% donkey serum and incubated overnight with a guinea pig polyclonal vGlut2 antibody (1:250, Millipore AB2251-1, RRID:AB_2665454). After primary antibody incubation, slices were rinsed 6×10 minutes and incubated with an AlexaFluor488-conjugated goat-anti-guinea pig IgG (1:200, Invitrogen A-11073, RRID:AB_2534117) for 4 hours, washed 3×5 min in PBS, and coverslipped with Vectashield Hardset. vGlut2 fluorescence was imaged on a Scientifica 2-photon microscope with a MaiTai HP Ti:sapphire laser tuned to 800 nm with a 370×370 micron field of view (2.77 px/micron). The signal in a single optical section was automatically thresholded and vGlut2 puncta with a size threshold of 6 μm^2^ were detected using the Synapse Counter plug-in in ImageJ [39]. For vGlut2 staining following brimonidine treatment, coronal sections containing the dLGN were stained using a rabbit polyclonal vGlut2 antibody (1:500, Cedar Lane/Synaptic Systems 135403(SY), RRID:AB_887883) and a donkey-anti-rabbit IgG secondary conjugated to AlexaFluor568 (1:200, Invitrogen A-10042, RRID:AB_2534017). Following staining, images were acquired with a 185×185 micron field of view (5.53 px/micron).

For measuring retinal ganglion cell density, retinas from D2 and D2-control mice (11-12 months age) were dissected free from the eyecup in Ames solution (US Biologicals, A13722525L). Relieving cuts were made and retinas were mounted on nitrocellulose membranes, after which they were fixed in 4% PFA for 30 mins. After 3×5 minute washes in PBS, retinas were blocked/permeabilized using a solution containing 1% TritonX-100, 5.5% donkey serum, 5.5% goat serum, and 0.5% dimethylsulfoxide for 1 hour. Following blocking and permeabilization, retinas were incubated overnight at 4°C in the same solution containing a rabbit-anti-choline acetyltransferase (ChAT) monoclonal antibody (1:1000, Abcam ab178850, RRID:AB_2721842) and a guinea pig-anti-NeuN polyclonal antibody (1:500, Millipore ABN90, RRID:AB_11205592). After 6×10 minutes of washing in PBS, retinas were incubated in AlexaFluor-conjugated secondary antibodies (1:200 goat-anti-guinea pig IgG 568, Invitrogen A-11075, RRID:AB_141954; 1:200 donkey-anti-rabbit IgG-488, Invitrogen A-21206, RRID:AB_2535792) for 4 hours, washed 3×5 mins, mounted on SuperFrost Plus slides, and coverslipped with Vectashield Hardset. NeuN and ChAT-labeled cells in the ganglion cell layer were imaged on a 2-photon microscope in 3-4 quadrants of the central retina (^~^500 microns from the optic nerve head) and peripheral retina (^~^1700 microns from the optic nerve head). RGCs counts were performed using an ImageJ macro in which the maximum intensity projections were thresholded, despeckled, and inverted followed by application of the “dilate”, “fill holes”, and “watershed” commands. Finally, the Analyze Particles tool was used to detect objects with a circularity of 0.3-1 with a size threshold of 30 μm^2^. Separately, ChAT-labeled amacrine cells were counted with a circularity of 0.4-1 and a size threshold of 80 μm^2^. In a subset of images (n=9), we found that 66% of ChAT^+^ cells detected using these parameters were also detected as NeuN^+^. Thus, the RGC number was taken as the difference of NeuN-labeled cells and NeuN/ChAT double-labeled cells, similar to the approach described previously [40]. Density was analyzed separately in central and peripheral retina using the mean cell counts analyzed in this manner from the 3-4 central or peripheral images for each eye.

### Patch-clamp electrophysiology

For measurements of retinogeniculate synaptic function, parasagittal sections containing the dLGN [41,42] were prepared using the “protected recovery” method [43,44] that involved sectioning in artificial cerebrospinal fluid (aCSF; 128 NaCl, 2.5 KCl, 1.25 NaH_2_PO_4_, 24 NaHCO_3_, 12.5 glucose, 2 CaCl_2_, and 2 MgSO_4_ and continuously bubbled with a mixture of 5% CO_2_ and 95% O_2_) followed by a 12-minute incubation in an N-methyl-D-glucamine-based solution (in mM: 92 NMDG, 2.5 KCl, 1.25 NaH_2_PO_4_, 25 glucose, 30 NaHCO_3_, 20 HEPES, 0.5 CaCl_2_, 10 MgSO_4_, 2 thiourea, 5 L-ascorbic acid, and 3 Na-pyruvate, warmed to 33°C). After an additional >1 hour recovery in aCSF at room temperature, slices were transferred to a recording chamber on an Olympus BX51-WI upright microscope and superfused with aCSF supplemented with 60 μM picrotoxin at ^~^2 mL/minute bubbled with 5% CO_2_ and 95% O_2_ and warmed to 30-33°C with an in-line solution heater (Warner Instruments).

Thalamocortical (TC) relay neurons were targeted for whole-cell voltage clamp recording using a pipette solution comprised of (in mM) 120 Cs-methanesulfonate, 2 EGTA, 10 HEPES, 8 TEA-Cl, 5 ATP-Mg, 0.5 GTP-Na_2_, 5 phosphocreatine-Na_2_, 2 QX-314 (pH = 7.4, 275 mOsm). Electrophysiology was performed using a Multiclamp 700B or 700A amplifier, a Digidata 1550B digitizer, and pClamp 10 or 11 software (Axon/Molecular Devices). The holding voltage was −70 mV after correction for the liquid junction potential, which was measured as 10 mV. A concentric bipolar stimulating electrode was positioned in the optic tract anterior and ventral to the ventral LGN (^~^1-1.5 mm from the dLGN) and used to deliver pairs of current stimuli to RGC axons from an AM Systems Model 2100 Isolated Pulse Stimulator (0.3-0.5 ms, 200 ms inter-stimulus interval).

To measure the maximal AMPA receptor-mediated excitatory post-synaptic currents (EPSCmax), representing the response evoked by all intact RGC inputs converging onto a given TC neuron [45–47], we increased stimulus intensity until the response amplitude plateaued (up to 10 mA stimulus amplitude). NMDA receptor EPSCs (EPSC_NMDA_) were measured as the amplitude 15 ms post-stimulus while the TC neurons were voltage clamped at +40 mV. To measure RGC convergence, the stimulus intensity was reduced until it evoked an EPSC representing the input from a single RGC axon (single-fiber EPSC, EPSCsf), defined as the EPSC amplitude recorded when a given stimulus failed to evoke a response approx. 50% of the time. The “fiber fraction” [45–47], which is an estimate of the single fiber contribution to the maximal EPSC, and thus, a metric for measuring RGC input convergence onto post-synaptic TC neurons, was calculated as the ratio of the EPSCmin/EPSCmax. These electrophysiology data were analyzed with ClampFit 11.

Single-vesicle miniature EPSCs (mEPSCs) were recorded over sixty seconds in the absence of stimulation. mEPSCs were detected and amplitude and frequency were analyzed using MiniAnalysis software (Synaptosoft, Fort Lee, NJ, USA) with an amplitude threshold of 4.5 pA. Fits of mEPSC amplitude and baseline noise histograms revealed good separation of mEPSCs from the noise with these detection parameters.

### Single-neuron dye fills and dendritic reconstruction

For single-neuron dendritic reconstructions, TC neurons were targeted for whole-cell patch clamp recording in 250 micron-thick coronal sections prepared using the protected recovery method, as described above. The pipette solution was either the Cs-methanesulfonate solution, as above, or was a K-gluconate solution (in mM, 120 K-gluconate, 8 KCl, 2 EGTA, 10 HEPES, 5 ATP-Mg, 0.5 GTP-Na_2_, 5 Phosphocreatine-Na_2_) supplemented with 2% Neurobiotin (Vector Laboratories, SP-1120) and 10 μM CF568 (Biotium). Neurobiotin and CF568 were injected using square-wave current injections (500 pA peak to trough, 2 Hz) for 10-15 minutes in current clamp mode, after which slices were fixed in 4% PFA for 1h and incubated for a week in 10 ug/mL streptavidin-568 in PBS with 1% TritonX-100 at 4°C. After incubation, slices were washed 3x 10 min in PBS, mounted on Superfrost Plus slides and coverslipped with Vectashield Hardset (Vector Laboratories, H-1400). Filled TC neurons were imaged on a 2-photon microscope and dendrites were reconstructed using Simple Neurite Tracer plug-in in ImageJ. Sholl analysis was performed in ImageJ on a 2-dimensional projection of the reconstructed dendrites (10 μm spacing between Sholl rings). Equivalent dendritic field diameter was calculated from the area of a convex polygon drawn by connecting TC neuron dendritic tips in ImageJ.

### Statistics

Statistical analysis was performed using GraphPad Prism 9. Normality of the data was assessed using a D’Agostino & Pearson test. When data were normally distributed, significance was assessed using a One-Way Analysis of Variance with Dunnett’s multiple comparison tests. To avoid pitfalls from pseudoreplication [48], statistical significance was measured using one-way nested ANOVA with a Dunnett’s multiple comparisons post-hoc test when we made multiple measurements from single animals (i.e., multiple cells from each mouse in a dataset). Nested data sets that followed a logarithmic distribution were log transformed prior to statistical testing. For all statistical tests, p<0.05 was considered statistically significant. Sample sizes (number of mice and the number of cells), statistical tests, and p-values are reported in the figure legends. Data are displayed as individual data points and mean ± standard error of the mean (SEM) or median ± inter-quartile range (IQR), as indicated in figure legends

## Results

To determine how IOP elevation in DBA/2J mice influences the dLGN, we performed experiments to longitudinally monitor IOP and assess anterograde axoplasmic transport of fluorescently-tagged cholera toxin beta (CTb) subunits (Figure 1). D2 mice showed an increase in IOP (n = 168 eyes, 84 mice) beginning at approximately 7 months of age. IOP elevation was variable, as we and others have reported previously. Compared to DBA/2J-gpnmb^+^ (D2-control mice, n = 110 eyes, 55 mice), IOP was significantly elevated after 8 months. Notably, IOP in D2 mice was significantly lower than in D2-controls at 5 months and 6 months (Figure 1 B&C), similar to what we have observed previously [49] and is apparent in figures from other prior studies [32,50,51]. While female DBA/2J mice had, on average, higher IOP than males at ages >8 months, consistent with prior work [32], the male and female IOP measurements overlapped considerably. Using serial histological sections of the dLGN, we quantified the total fraction of dLGN labeled by anterogradely-transported CTb-594 (Figure 1D&E). Approximately 80% of the dLGN was labeled in D2-control mice, which was similar to the amount labeled in 4 month-old D2 mice and was the result of complete labeling from contralateral projections and no labeling in the ipsilateral projection region. By 9 months of age, 29±1% of the dLGN was labeled, and the pattern appeared to be the result of regional loss of transport, similar to what has been documented for the superior colliculus. There was also a weak but statistically significant correlation of CTb-594 labeling with the summed IOP measurements taken for the two months prior to tissue collection (Figure 1F), suggesting a functional link between IOP and deficits in anterograde transport to the dLGN.

**Figure 1.**
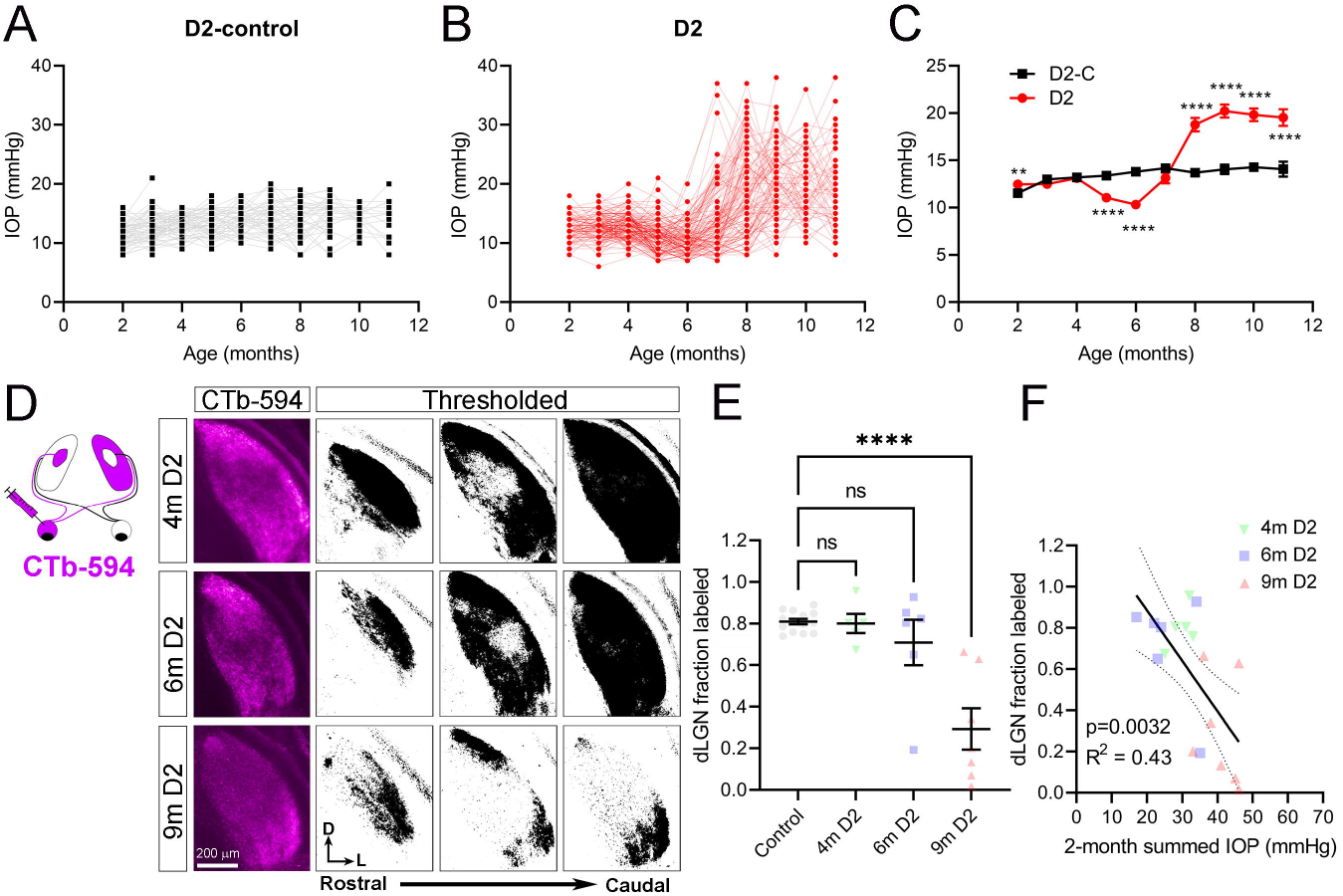
Elevated intraocular pressure and deficits in anterograde transport to the dLGN. **A)** Intraocular pressure (IOP) measurements from DBA/2J-gpnmb+ (D2-control) mice (n = 110 eyes from 55 mice included in this study). **B)** IOP measurements from DBA/2J (D2) mice (n = 168 eyes from 84 mice). **C)** Mean (± SEM) IOP measurements from D2 and D2-control eyes. Unpaired t-tests: 2m (month) p=0.0038; 3m p=0.13; 4m p=0.78; 5m p=1.0×10^−10^; 6m p=2.3×10^−14^; 7m p=0.14; 8m p=6.6×10^−9^; 9m p=4.0×10^−11^; 10m p=1.3×10^−9^; 11m p=2.5×10^−5^. **D)** Fluorescently-tagged cholera toxin-beta (CTb) was injected unilaterally and the area of labeled contralateral dLGN was measured based on fluorescence signal in serial dLGN sections. **E)** Group data (mean ± SEM) showing fraction of CTb-labeled dLGN. There was a significant difference among groups (one-way ANOVA, p=5.2×10^−6^) and the 12m group significantly differed from the control group (Dunnett’s multiple comparison test; p<1×10^−15^). **E)** For the D2 mice, there was a significant negative correlation (linear regression with 95% confidence interval) of the fraction of dLGN labeled by CTb with the sum of the two IOP measurements taken prior to tissue collection (Pearson correlation, p=0.0032).

To test the influence of age and IOP on RGC axon terminals, we next immunostained dLGN sections for vGlut2 (Figure 2). In these experiments, the density of vGlut2-labeled puncta was comparable between D2-control and 4 month-old D2 mice. In older D2 mice, the density of vGlut2-labeled puncta showed a progressive reduction, being lower in 9m and lower still in 12m D2 mice. To better understand the relationship between loss of vGlut2 puncta and deficits in anterograde transport to the dLGN, we performed vGlut2 immunofluorescence staining on dLGN sections that had also been labeled via anterograde CTb transport (Figure 2C-F). We compared vGlut2 density in regions of 9m D2 dLGN with intact CTb labeling (“CTb-intact”) with dLGN regions that had little CTb labeling (“CTb-deficient”). In this experiment, we found that CTb-deficient regions of the dLGN also had lower vGlut2 puncta density compared to vGlut2 density in the CTb-intact regions. However, this was not a one-to-one relationship; while there was a correlation of CTb labeling intensity with vGlut2 puncta density, regions with very little CTb still had labeling for vGlut2. This suggests that loss of vGlut2 signal could be due to either diminished vGlut2 transport from the dLGN or due to degeneration of RGC axon terminals.

**Figure 2.**
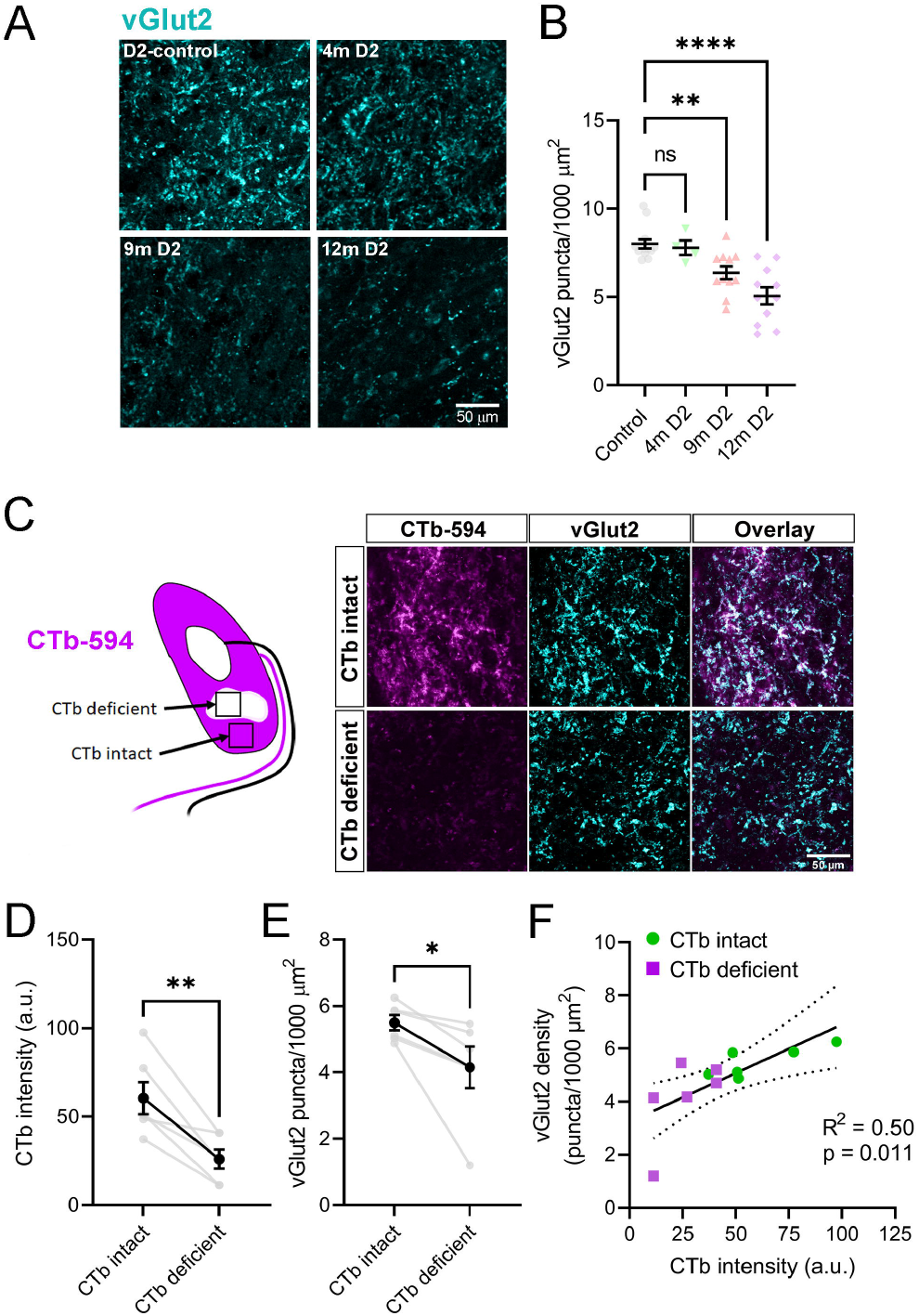
Loss of vGlut2 labeled RGC axon terminals in the dLGN is associated with transport deficits in DBA/2J mice. **A)** Single optical sections of dLGN labeled with an anti-vGlut2 antibody from a D2-control mouse and D2 mice at 4m, 9m, and 12m of age. **B)** Group data (mean ± SEM) showing density of detected vGlut2 puncta. There was a significant difference among groups (one-way ANOVA, p=8.0×10^−6^) with 9m and 12m groups differing significantly from the control group (Dunnett’s multiple comparison: 4m p=0.98; 9m p=0.0070; 12m p<1×10^−15^). **C)** Analysis of vGlut2 density in regions of the dLGN with intact or deficient anterograde transport of unilaterally-injected CTb. **D)** Quantification (mean ± SEM) of CTb pixel intensity in “intact” or “deficient” dLGN regions (p=0.0065, paired t-test). **E)** Quantification of vGlut2 density (mean±SEM) in dLGN regions with intact or deficient CTb labeling (p=0.041, paired t-test). F) Significant positive correlation (linear regression with 95% confidence interval) of vGlut2 density with intensity of CTb labeling (Pearson correlation, p=0.011).

The above data imply that impaired axonal function (as indicated by reduction in anterograde transport) is related to an impairment of a presynaptic structural marker of RG synapses (vGlut2). Therefore, we next sought to determine the consequences of elevated IOP on RG synaptic function by performing whole-cell voltage-clamp recordings of dLGN TC neurons in a parasagittal slice preparation. First, we recorded mEPSCs in the absence of stimulation in D2-control and D2 mice at 6, 9, and 12 months of age (Figure 3). We found that mEPSC amplitude was not significantly different between groups, suggesting that there was no detectable alteration in AMPA receptor properties or composition at RG synapses. However, there was a statistically significant difference in mEPSC frequency between groups. While it was similar in D2-controls and 6m D2 mice, there was a significant reduction in mEPSC frequency compared to controls at 9m and 12m of age, similar to what we have shown in coronal sections in D2 mice and mice with IOP elevation from anterior chamber microbead injections.

**Figure 3.**
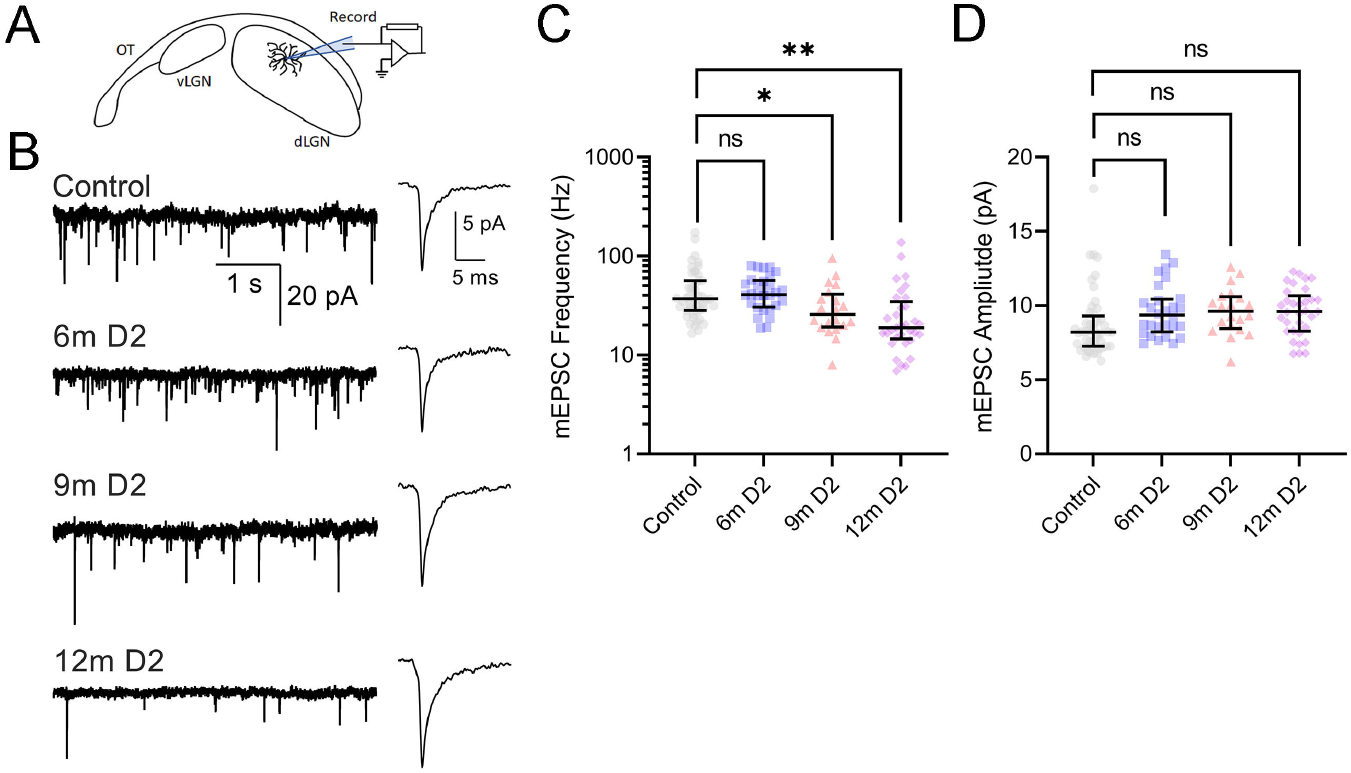
Progressive loss of miniature excitatory post-synaptic currents recorded from dLGN thalamocortical relay neurons in DBA/2J mice. **A)** Recording schematic of optic tract (OT), ventral lateral geniculate nucleus (vLGN), and dLGN with patch clamp electrode in parasagittal slice. **B)** Left: Example 5-second duration traces of miniature excitatory post-synaptic currents (mEPSCs) recorded in the absence of stimulation from D2-control and D2 mice. Right: average of the detected mEPSC waveforms. **C)** Group data (median ± IQR) of mEPSC frequency. There was a significant difference among groups (nested one-way ANOVA, p=0.0021) and 9-month and 12-month groups differed significantly from control (Dunnett’s multiple comparison: 6m p=0.99; 9m p=0.026; 12m p=0.0078). **D)** Group data (median ± IQR) of mEPSC amplitude. There was no significant difference among groups (nested one-way ANOVA, p=0.19) and individual groups not significantly different from the control (Dunnett’s multiple comparison, 6m p=0.24; 9m p=0.29; 12m p=0.21). Group sizes: control n=48 cells, 13 mice; 6m n=28 cells, 7 mice; 9m n=20 cells, 8 mice; 12m n=33 cells, 10 mice.

A reduction in mEPSC frequency can be attributed to a reduction in the number of functional synapses and/or a reduction in the probability of vesicle release. To probe these possibilities, we used a stimulating electrode positioned in the optic tract to stimulate RGC axons while we recorded the AMPA receptor-mediated EPSCmax (Figure 4), which represents the contributions from all intact axons converging onto a given TC neuron. In these experiments, we found that there was a statistically significant difference in EPSCmax between groups, with a detectable reduction in EPSCmax in slices from 12 month-old D2 mice compared to controls (Figure 4A-C). There was a weak but statistically significant correlation of Log10(EPSCmax) (averaged across cells in each mouse) with the cumulative IOP measurements taken prior to conducting electrophysiology experiments (R^2^=0.23, p=0.013). The lower EPSC amplitude was mirrored when we recorded NMDA receptor-mediated EPSCs (EPSC_NMDA_) by changing the holding potential to +40 mV (Figure 4D). There was no significant change in the AMPA/NMDA (Figure 4E) ratio suggesting no changes in the relative contributions of each receptor type at RG synapses or “silent synapses” contributing to the reduced EPSC amplitude. When we delivered a pair of stimuli to the optic tract (200 ms inter-stimulus interval), we found that there was a statistically significant difference among the groups, with an increase in the ratio of the second response to the first (Paired pulse ratio/PPR; EPSC2/EPSC1) detectable in 12m D2 mice compared to controls (Figure 4F). In some cases, the PPR was >1, which represents a shift from the synaptic depression more typical for RG synapses to a mode of synaptic facilitation, suggestive of a decrease in presynaptic vesicle release probability.

**Figure 4.**
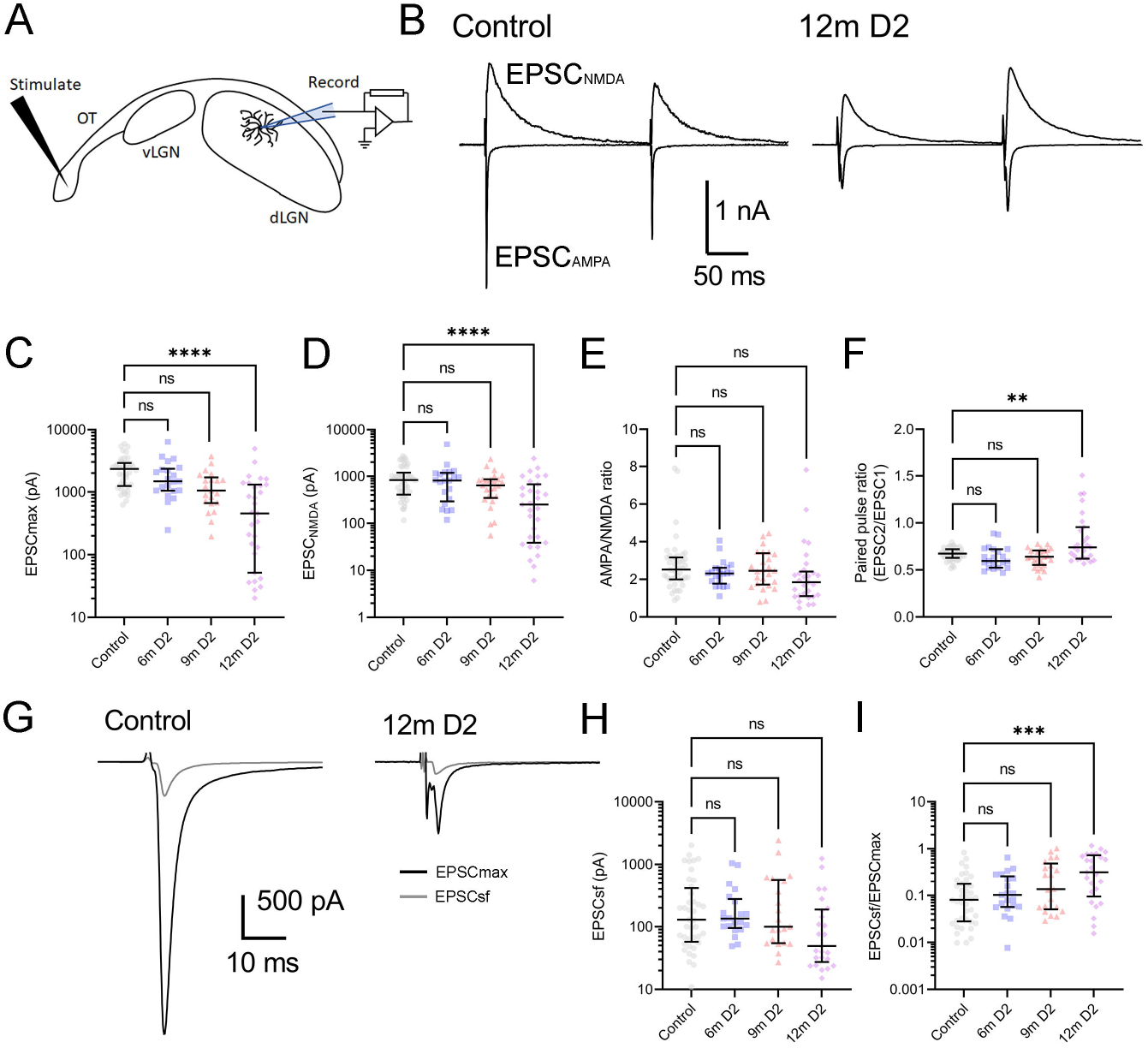
Progressive loss of convergent retinal inputs to dLGN relay neurons in DBA/2J mice. **A)** Recording schematic of optic tract (OT) with stimulating electrode, ventral lateral geniculate nucleus (vLGN), and dLGN with patch clamp electrode in parasagittal slice. **B)** Example maximal AMPA- and NMDA-receptor-mediated EPSCs recorded at −70 mV and +40 mV, respectively, following maximal stimulation of the optic tract with a pair of pulses (200 ms inter-stimulus-interval). **C)** Group data of the AMPA-receptor-mediated EPSCmax shows that the EPSC differed among the groups (nested one-way ANOVA, p= 2.4×10^−5^). The EPSCmax was significantly smaller amplitude in recordings from 12m D2 mice compared to controls (Dunnett’s multiple comparison test: 6m p=0.77; 9m p=0.16; 12m p<1×10^−15^). **D)** The NMDA-receptor-mediated EPSC differed among groups (nested one-way ANOVA, p=3.6×10^−5^) and the 12m amplitudes were significantly lower than control (Dunnett’s multiple comparison: 6m p=0.97; 9m p=0.54; 12m p<1×10^−15^). **E)** The AMPA/NMDA ratio did not significantly differ across groups (nested one-way ANOVA, p=0.34). **F)** Paired pulse ratio differed among groups (nested one-way ANOVA, p=0.0011) and was significantly higher in 12m D2 mice compared to controls (Dunnett’s multiple comparison: 6m p=0.75; 9m p=0.79; 12m p=0.0040). C-F) Show median ± IQR. Sample sizes: Control, n=40-46 cells, 12-14 mice; 6m n=21 cells, 7 mice; 9m n=21 cells, 9 mice; 12m n=29-31 cells, 10 mice. **G)** Example maximal EPSCs and single-fiber EPSCs (EPSCsf) from a control and 12m D2 mouse. **H)** The single-fiber EPSC amplitude (median ± IQR) did not differ among groups (nested one-way ANOVA, p=0.49). **I)** The “fiber fraction” (EPSCsf/EPSCmax; median ± IQR) differed among groups (nested one-way ANOVA, p=0.0038) and the 12m value was significantly different from control (Dunnett’s multiple comparison; 6m p=0.99, 9m p=0.16; 12m p=0.0022).

When we reduced the stimulus amplitude to evoke synaptic vesicle release from a single RGC axon, we found that there was no significant difference in the EPSCsf among groups (Figure 4G&H). The ratio of EPSCsf/EPSCmax (the “fiber fraction”) has been used to monitor the developmental refinement of RGC inputs onto TC neurons and represents a statistically quantifiable estimate of synaptic convergence [45–47]. We found that EPSCsf/EPSCmax was significantly increased in 12m D2 mice compared to controls, indicating a reduction in the number of functional RGC axon inputs onto each TC neuron.

Taking the reciprocal of the fiber fraction suggests that each TC neuron receives inputs from an average 6.9 RGC axons in control mice, while 12m D2 mice receive an average of 2.3 functional RGC axon inputs. At the same time point, we detected a reduction in mEPSC frequency by 18 Hz; control mEPSC frequency was 37 Hz while it was 19 Hz in 12m D2 mice. Comparison with published anatomical studies of RG synapses points to the congruence of these two measurements [52,53]; if each RGC axon contributes 15 boutons to a post-synaptic TC neuron, each with 27 active zones having a spontaneous vesicle fusion rate of 0.01 Hz per active zone [54], then the expected result of a drop from 6.9 RGC axonal inputs to 2.3 inputs is an 18 Hz reduction in mEPSC frequency. This indicates that the change in synaptic transmission measured with both mEPSCs and optic tract stimulation are of a similar scale and likely to represent complementary measures of a similar pathological process.

The above findings show a loss of vGlut2-labeling of RGC axon terminals and diminished numbers of RGC inputs to each TC neuron in older DBA/2J mice. However, post-synaptic TC neuron structure and function are likely to be altered in glaucoma as well. For instance, elevated IOP is associated with altered TC neuron intrinsic excitability and somatic atrophy. Prior studies have also reported reorganization of LGN neuron dendrites in late-stage glaucoma in primates [27,55,56] and we have previously shown changes in TC dendritic structure following bilateral enucleation [26], a traumatic form of optic nerve injury, and IOP elevation with microbead injections [14]. Here, we performed Sholl analysis of TC neuron dendrites reconstructed after neurobiotin filling during whole-cell recording and found that although TC neuron dendritic complexity was comparable between control and 9m D2 mice, there was a modest reduction in the peak number of Sholl intersections in 12m D2 mice compared to controls (Figure 5). There was no statistically significant difference in dendritic field diameter among the groups. Thus, reorganization of post-synaptic TC neuron structure accompanies loss of function RG synaptic inputs.

**Figure 5.**
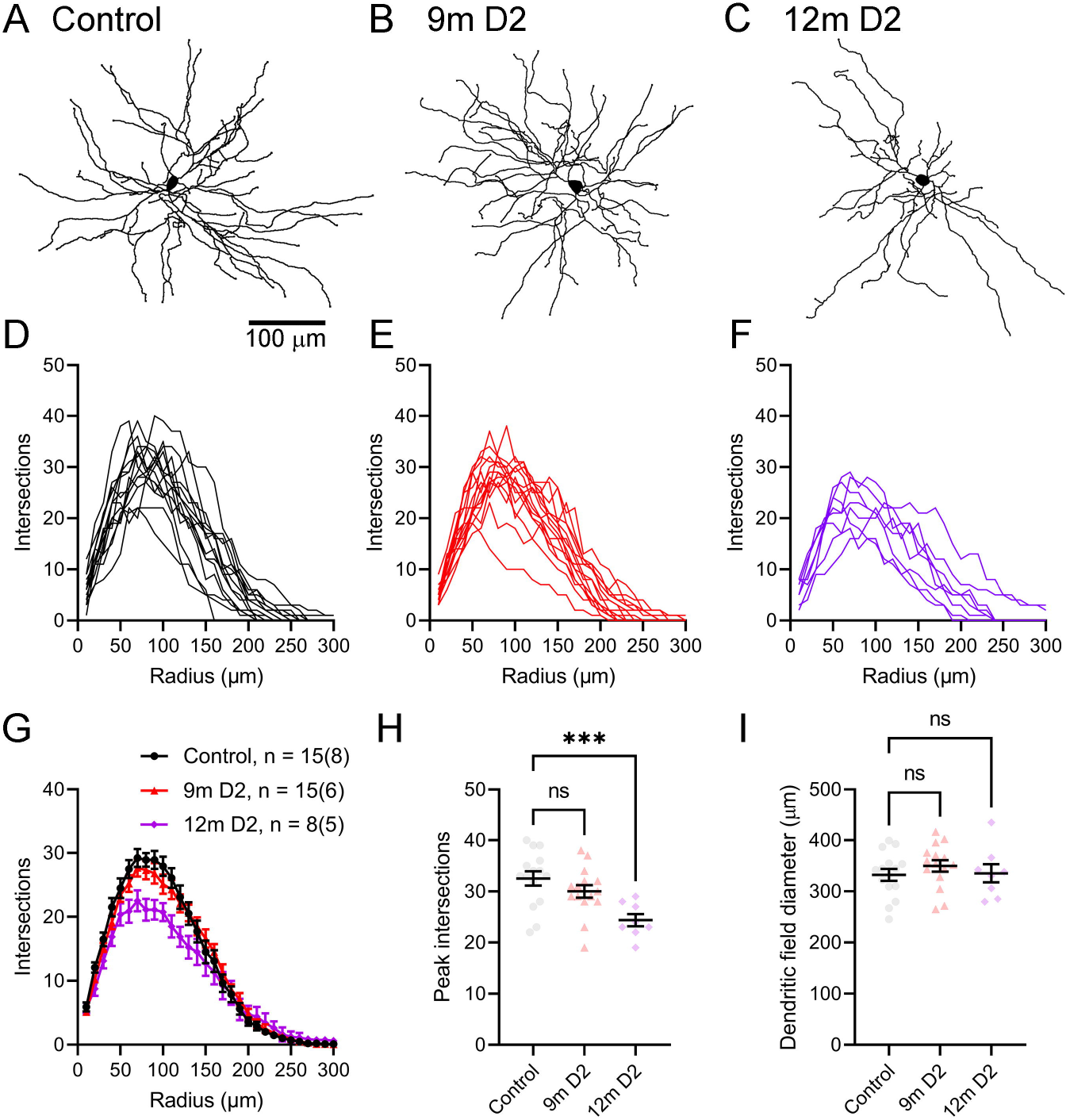
Thalamocortical neuron dendritic remodeling in DBA/2J mice. **A-C)** Reconstructed TC neurons from D2-control (A), 9m D2 (B), and 12m D2 (C) mice filled with Neurobiotin during whole-cell recording in coronal slices. **D-F)** Sholl plots of each TC neuron included in sample. **G)** Group data (mean ± SEM) of Sholl plots. **H)** Group data (mean ± SEM) of the peak number of Sholl intersections for each cell. There was a significant difference among groups (nested one-way ANOVA, p=0.0033) and the 12m peak intersections was significantly lower than control (Dunnett’s multiple comparison, 9m p=0.46; 12m p=0.0019). **I)** Group data (mean ± SEM) of the dendritic field diameter measured as the equivalent diameter of a convex polygon of the dendritic field. There was no statistically significant difference among groups (nested one-way ANOVA, p=0.80). Sample size: Control n = 15 cells (8 mice); 9m D2 n=15 cells (7 mice); 12m D2, n=8 cells (5 mice).

We next sought to count the number of RGC somata in DBA/2J retinas to provide another comparison of the above findings with a commonly used metric of glaucomatous progression (Figure 6). Some commonly used markers for RGC somata such as RBPMS or Brn3 appear to undergo expression changes in DBA/2J retinas, meaning that using them to measure RGC somatic degeneration might lead to undercounts of RGC density and consequent over-estimation of RGC somatic degeneration [40,57–59]. The use of the neuronal marker NeuN, in conjunction with correction of counts for NeuN-positive cholinergic amacrine cells in the ganglion cell layer (identified by labeling for choline acetyltransferase; ChAT) is a reliable way of counting RGCs and has been used previously to show that somatic loss is a late event in DBA/2J mice [2,40]. We found that the density of ChAT^+^ cells in the RGC layer was not significantly different between D2 and D2-control mice (11-12 months age), although it was slightly lower than previously reported in C57Bl/6J and A/J mice, which likely reflects strain differences [60,61]. We measured the density of NeuN^+^ presumptive RGCs by correcting for ChAT^+^/NeuN^+^ double-labeled cells, finding that there was no significant difference in the density of NeuN^+^ RGCs. This is consistent with prior work suggesting that RGC somatic loss is a late event in DBA/2J mice [40].

**Figure 6.**
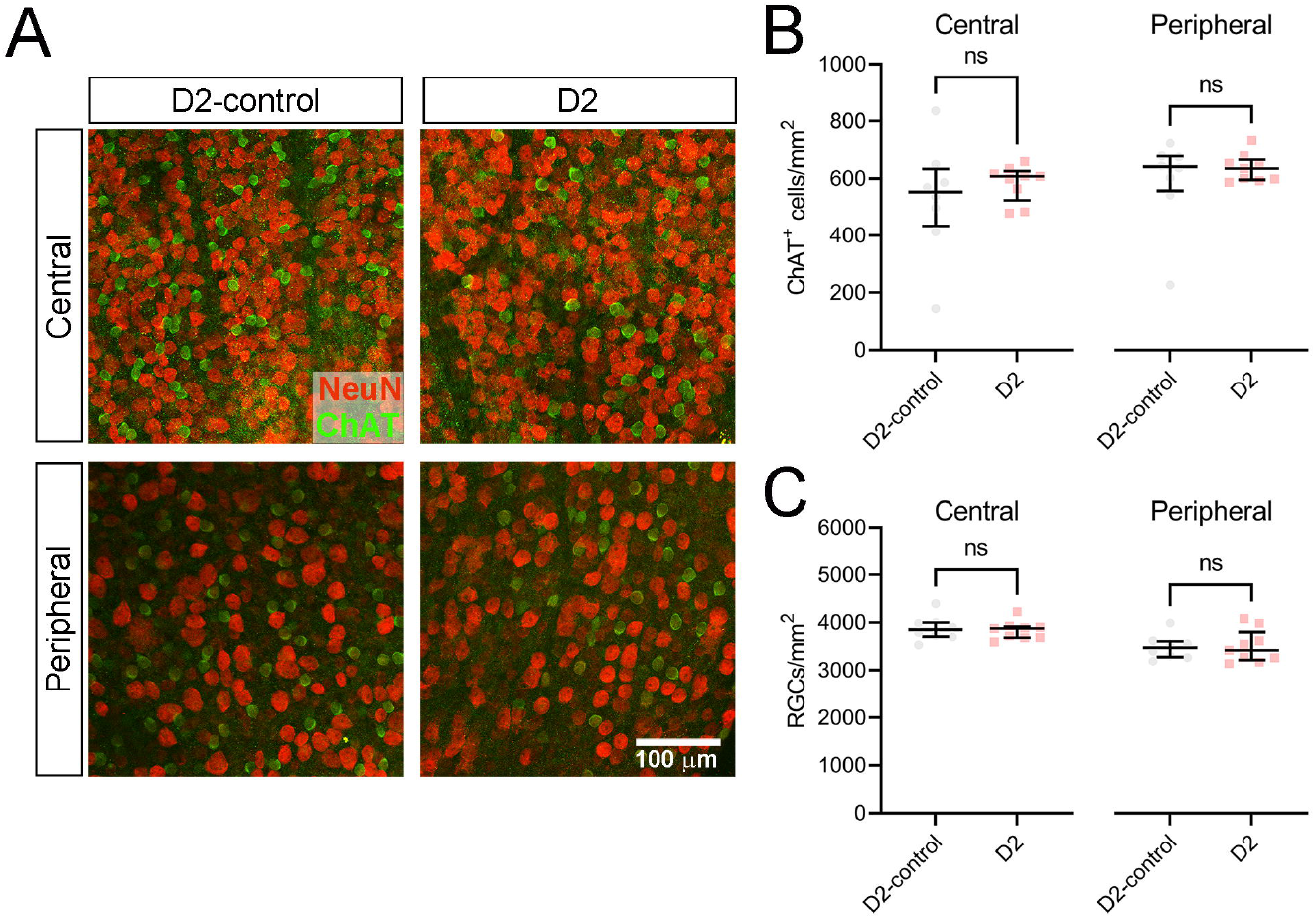
No loss of retinal ganglion cell somata in 11-12-month-old DBA/2J mice. **A)** 2-photon Immunofluorescence images of retinal flat mounts from a 12-month-old D2-control mouse and a 12 month-old D2 mouse stained with antibodies for NeuN and choline acetyl transferase (ChAT). Images were acquired from central retina (centered ^~^500 microns from the optic nerve head) and peripheral retina (centered ^~^1700 microns from the optic nerve head). **B)** Analysis of ChAT^+^ cell density (median ± IQR). Each data point is the ChAT^+^ cell density averaged across three to four quadrants for each retina. There was no significant difference in ChAT^+^ cell density between D2-control and D2 mice (central p=0.44; peripheral p=0.42, unpaired t-test). **C)** RGC density was measured as the difference between the total number of NeuN^+^ cells and the number of NeuN^+^/ChAT^+^ double-labeled cells. There was no significant difference between D2-control and D2 RGC density (central p=0.68; peripheral p=0.93, unpaired t-test). Sample sizes: D2-control, n=8 retinas, 4 mice; D2, n=9 retinas, 5 mice.

Finally, we tested whether treatment with IOP-lowering medication could prevent effects on dLGN synaptic structure in D2 mice (Figure 7). To do this, we treated a subset of DBA/2J mice with brimonidine eye drops daily for 4-5 days per week from approximately 6 months to 9 months of age prior to harvesting tissue. IOP measurements in 7-9-month old DBA/2J mice showed treatment was followed by an acute 5.6+/-1.4 mmHg reduction in IOP (Figure 7C; n = 20 eyes, 10 mice; p = 0.0011). Brimonidine has been shown to reduce IOP in mice over a 24-hour period [62] but drug effects do wash out over time. Indeed, IOP values taken earlier in the day, prior to brimonidine administration were similar in brimonidine-treated eyes compared to those receiving only lubricating eye drops and this was evidenced by similar IOP values at the 8-8.5m time point (23.2±2.3 mmHg for lubricating eye drops group and 22.8±1.7 for brimonidine group; p=0.88). Ultimately, treatment with brimonidine reduced the loss of vGlut2-labeled axon terminals (Figure 7D-F); we found vGlut2 density was higher in dLGN from brimonidine treated mice relative to those receiving only lubricating eye drops (p=0.0041).

**Figure 7.**
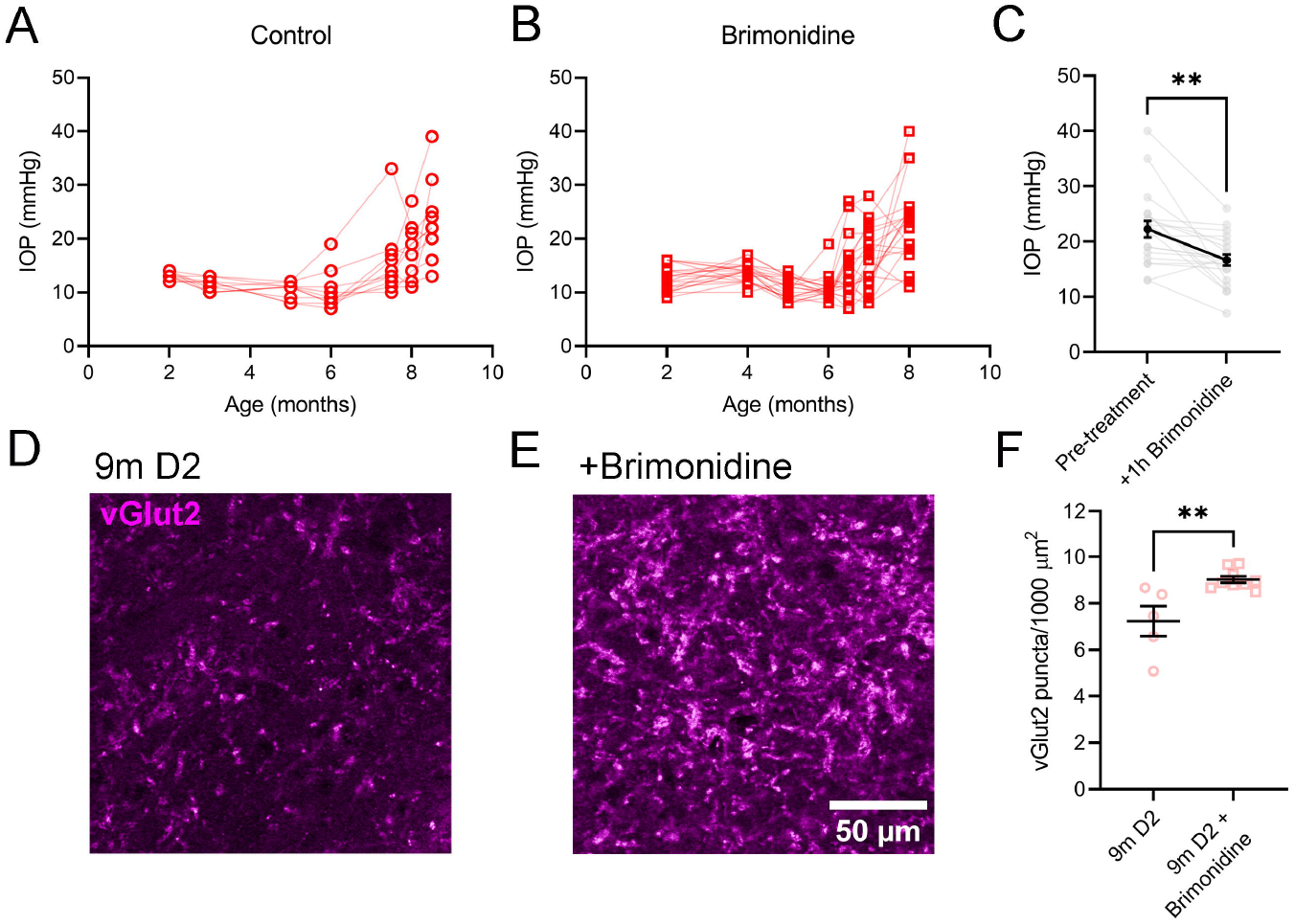
Treatment with brimonidine eye drops saves vGlut2 labeling in DBA/2J mice. **A, B)** IOP measurements from D2 mice treated (4-5 days/week) with either lubricating eye drops **(A)** or 0.2% brimonidine tartrate eye drops **(B)** taken ^~^20h following previous eye treatment. Treatment started at approx. 6m of age and lasted through 9m of age. Both groups had similar IOP profiles and 8-8.5m IOP was similar between groups (p=0.87, unpaired t-test). **C)** IOP measurements before and 1h post-brimonidine treatment showing that brimonidine eye drops acutely lowered IOP (p=0.0011, paired t-test). **D, E)** Single 2-photon optical sections of vGlut2 immunofluorescence labeling of vGlut2 in dLGN of D2 mouse treated with lubricating eye drops (**D**, “9m D2”) or a mouse treated with brimonidine (**E**, “+Brimonidine”). **F)** Quantification of vGlut2 puncta (mean ± SEM) showing a higher density of RGC axon terminals in mice treated with brimonidine (p=0.0041, unpaired t-test). Sample sizes: control (lubricating eye drops group), n = 10 eyes, 5 mice; brimonidine group, n = 18 eyes, 9 mice.

## Discussion

The results here demonstrate a loss of retinal ganglion cell output synapses in the dLGN in a mouse model of glaucoma occurring prior to degeneration of RGC somata. This appears to involve the drop-off of individual RGC axon inputs to post-synaptic TC neurons without appreciable alterations in the strength of individual RGC synapses. Notably, we find that IOP elevation is associated with a diminishment of anterograde optic tract transport to the dLGN and loss of vGlut2-labeling of RGC synaptic terminals. The transport deficits were associated with diminished vGlut2 labeling, although the loss of transport was more severe than the loss of vGlut2, suggesting that compromised axonal function precedes the loss of vGlut2 labeling. These pre-synaptic effects were followed by modest loss of dendritic complexity in proximal regions of TC neuron dendritic arbors, which might represent a disruption of dendritic homeostasis due to diminished retinogeniculate synaptic inputs. Finally, treatment with brimonidine eye drops over a period of three months beginning slightly prior to eye pressure elevation in D2 mice saved the vGlut2 labeling, highlighting the potential for therapeutic interventions to modulate synaptic dysfunction in the dLGN.

In DBA/2J mice, we found that mEPSC frequency recorded from TC neurons was reduced, which is consistent with what we have found previously using coronal slices from D2 mice or mice with experimentally-elevated IOP [14,25]. The use of fiber fraction measurements of RGC convergence in control mice were consistent with values obtained previously [45–47]. These measurements showed that TC neurons from 12m D2 mice receive inputs from fewer RGCs than observed in controls. While the mEPSC results would be consistent with several scenarios including loss of synaptic contacts made by each RGC axon or weakening of individual inputs, our data obtained with optic tract stimulation do not indicate a change in the number of bouton contacts or a major change in the strength of individual synapses, as either process would be reflected in a reduction in the EPSCsf, which we did not find. This instead largely results from drop-off of individual RGC axons, as evidenced by the increase in fiber fraction and no statistically detectable change in single fiber strength.

Our vGlut2 labeling studies likewise align with this picture; namely, they show a diminishment of vGlut2 labeling apparent at 9 months and to a greater extent at 12 months. vGlut2 is thought to be a relatively specific marker of RGC axon terminals in the dLGN as vGlut2 labeling is generally lost following enucleation [26,63–67]. We found that vGlut2 density was related to the extent of anterograde transport in dLGN regions with intact or deficient CTb labeling but that a considerable amount of vGlut2 was still present even in regions with minimal CTb. This is consistent with the notion that loss of RGC axon terminal labeling lags transport deficits in the dLGN. Studies of the superior colliculus, another major RGC projection target, showed pronounced CTb transport labeling deficits despite intact vGlut2 labeling in D2 mice [17].

We hypothesize that loss of vGlut2 labeling results from deficient anterograde transport, albeit with a delay relative to CTb transport deficits, and that the terminals themselves remain even after total loss of vGlut2 immunopositivity. Synaptic proteins are either translated in the soma and actively transported to presynaptic terminals or are produced as the result of local axon terminal translation from mRNAs transported from the soma [68]. Presynaptic proteins have a slow turnover relative to other cellular proteins [69,70] and the pace of this process, in concert with intact local translation, could account for the persistence of vGlut2 labeling in regions deficient in anterograde transport.

Ultrastructural studies of the D2 superior colliculus (SC) show that RGC axon terminals persist in regions deficient for transported CTb [17] although they have altered ultrastructural properties including atrophied terminal size, misshapen mitochondria, and smaller active zones [24]. This might reflect loss of synaptic function despite the structural persistence of the terminals. Moreover, vGlut2 is important for loading glutamate into presynaptic vesicles, so loss of vGlut2 would lead to compromised synaptic function. vGlut2 expression also appears to regulate vesicle release probability [71], which might contribute to the changes in PPR. Notably, we did not detect any statistically significant differences in vGlut2 punctum size between transport intact and deficient regions at the level of light microscopy in the dLGN. While this might represent a contrast with the RGC axon terminal pathology in the SC [24], future ultrastructural studies along measurements of mitochondrial function will be necessary to test this in the dLGN. If the effects on RGC terminals in the SC apply to those in the dLGN, deficits in anterograde transport likely correspond to compromised axonal health and consequent loss of synaptic function, although we have not yet tested this possibility.

In addition to the presynaptic deficits (diminished vGlut2 labeling and anterograde transport, drop-off of RGC axon synaptic output, etc.), we also show that TC neurons in aged D2 mice display reorganization of their post-synaptic dendrites. Such dendritic reorganization is a common feature of neurodegenerative diseases [72]. Prior evidence from primate LGN has identified some dendritic loss in glaucoma [27,55,56] and we have previously found reductions in TC neuron dendritic complexity following experimental IOP elevation with microbeads and following bilateral enucleation [14,26]. Retinal input is important for organization of TC neuron dendrites, as evidenced by studies with mice that fail to develop retinogeniculate projections [73,74]. Here, in 12m D2 mice, we find reduced complexity in regions of TC neuron dendritic arbors proximal to the soma. This region of the TC neuron dendrites has a higher concentration of retinogeniculate “driver” inputs compared to the distal dendrites [53,75], where there is a greater concentration of “modulator” excitatory inputs arising from corticothalamic feedback synapses. Synaptic inputs are important for dendritic maintenance, with deafferentation or reduced synaptic strength being salient triggers for dendritic loss [72,76,77]. This is a potentially important role of spontaneous synaptic transmission, with mEPSCs serving a homeostatic role for synaptic maintenance. Thus, it is likely that dendritic loss here is primarily a response to rather than a cause of diminished retinogeniculate synaptic strength. It remains to be tested, however, whether the loss of postsynaptic dendritic complexity is preceded by loss of postsynaptic contacts (i.e. PSD-95 puncta) in D2 TC neurons.

Finally, what is the relationship between dLGN synaptic function and RGC somatic degeneration? Vision impairment in glaucoma is sometimes linked with RGC apoptosis, although numerous degenerative events in each of the RGC “compartments” - somatic/dendritic, optic tract, and axon terminal – precede detectable somatic loss. Additionally, measurements of RGC somatic loss are often complicated by challenges arising from labeling approaches and no labeling method is without drawbacks. For instance, commonly used RGC-labeling antibody markers such as RBPMS or Brn3 appear to have reduced expression in D2 retinas [57–59], which might lead to undercounting RGCs and over-estimating degeneration. Likewise, identification of RGCs via retrograde labeling can be confounded by optic nerve transport deficits [78], leading to similar over-estimations of degeneration. Here, we used an immunofluorescence approach that has been employed previously to count RGCs in D2 mice, which showed that that RGC loss in D2 mice is a late event, not occurring until after approximately 15 months of age. Consistent with this, we found no detectable RGC loss in 11-12-month-old D2 mice in our colony, indicating that loss of RGC output synaptic function in the dLGN occurs prior to loss of RGC somata in the retina. While our results indicate that loss of dLGN vGlut2 labeling and RGC synaptic function occurs prior to RGC somatic degeneration, RGCs undergo numerous other structural and functional changes prior to somatic loss in D2 mice. These include altered dendritic complexity and synapse loss [5–7,9,10,26], enhanced intrinsic excitability [5–7], altered light responses [5–7], disrupted metabolic function [21,79], optic nerve atrophy and gliosis, etc. Thus, the results of the current study support the body of evidence indicating that pathological changes to visual system structure and function prior to RGC somatic loss – in this case, diminishment of visual information transfer at the retinogeniculate synapse – contribute to visual impairment in glaucoma.

### Limitations and future directions

There are several limitations with the current study and areas for future exploration. First, electrophysiology studies were conducted in the presence of picrotoxin to isolate excitatory inputs from feed-forward or feedback inhibitory circuits in the dLGN. Consequences of glaucoma on dLGN inhibitory circuits remain to be explored and understanding the nature of such influences will be important for a more complete picture of dLGN function. Second, as discussed above, these results do not differentiate whether glaucoma leads to degenerative loss of synapses at this time point, as loss of vGlut2 labeling likely results from deficits in axon transport. Moreover, while we show that TC neurons lose post-synaptic dendritic complexity in 12m D2 mice, we do not know whether this is preceded by a loss of post-synaptic sites (such as PSD95 puncta) that might contribute to the diminished synaptic strength. Ultrastructural studies of both pre- and post-synaptic contacts will be needed to ascertain whether these structures remain intact. Third, differences in dLGN regions receiving input from RGC axons with intact vs. deficient CTb transport might contribute to the variability in measurements of synaptic function we show here. Our data point to a relationship between axon transport integrity and vGlut2 and prior work has shown that transport integrity deficits are related to defects in ultrastructure of presynaptic RGC axon terminals in the SC [17,24,78]. We have not yet explored the link between transport integrity and synaptic function in the dLGN. Fourth, while we show that treatment with brimonidine saves RGC axon terminals, we do not yet know whether this is due to brimonidine reducing IOP, to its neuroprotective effects, or to both [34–37,80–82]. We also have not yet explored the consequences of brimonidine treatment on other parameters such as electrophysiological measures of synaptic function or anterograde transport, although Lambert et al. have shown previously that systemic brimonidine administration prevents anterograde transport deficits and other changes in RGC morphology and optic nerve function [34]. Additionally, our experimental design with brimonidine involved treatment beginning prior to IOP elevation, so it will be important to determine if such treatments can reverse pathology if started at a later time point. Finally, while we have shown previously that TC neurons become more excitable in glaucoma [25], which might represent a homeostatic attempt to maintain thalamocortical information transfer following diminished retinogeniculate synaptic strength, we have not yet explored the consequences of these two phenomena operating in concert; it is possible that enhanced TC neuron excitability serves to maintain signaling to the visual cortex until a tipping point in the disease process after which the fidelity of visual signaling is impaired.

## Declarations

### Ethics approval and consent to participate

Animal studies were approved by the Institutional Animal Care and Use Committee at the University of Nebraska Medical Center.

### Consent for publication

Not applicable

### Availability of data and materials

The datasets used and/or analysed during the current study are available from the corresponding author on reasonable request.

### Authors’ contributions

All authors were involved in collecting and analyzing data. MJVH drafted the manuscript and prepared figures. All authors read and approved the final manuscript.

## Acknowledgements

Not applicable.

